# Application of components analysis in dynamic thermal breast imaging to identify pathophysiologic mechanisms of heat transfer

**DOI:** 10.1101/2021.08.16.456538

**Authors:** Meir Gershenson, Jonathan Gershenson

## Abstract

**Significance:** Of the most compelling unsolved issues in the paradigm to create success in the field of breast cancer infrared imaging is localization of direct internal heat of the tumor. The contribution of differential heat production related to metabolism versus perfusion is not understood. Previous work until now has not shown progress beyond identifying veins which are fed by the ‘hot’ cancer. Employing signal analysis techniques, we probe important questions which may lead to further understanding pathophysiology of heat transfer occurring in the setting of malignancy.

**Aim:** When using thermal imaging to detect breast cancer, the dominant heat signature is that of indirect heat transported in gradient away from the tumor location. Unprocessed images strikingly display vasculature which acts to direct excess heat superficially towards the skin surface before dissipating. In current clinical use, interpretation of thermogram images considers abnormal vascular patterns and overall temperature as indicators of disease. The goal of this work is to present a processing method for dynamic external stimulus thermogram images to isolate and separate the indirect vascular heat while revealing the desired direct heat from the tumor.

**Approach:** In dynamic thermal imaging of the breast, a timed series of images are taken following application of external temperature stimulus (most often cooled air). While the tumor heat response is thought to be independent of the external stimulus, the secondary heat of the veins is known to be affected by vasomodulation. The recorded data is analyzed using independent component analysis (ICA) and principal component analysis (PCA) methods. ICA separates the image sequence into new independent images having a common characteristic time behavior. Resulting individual components are analyzed for correspondence to the presence or lack of vasomodulation.

**Results:** Using the Brazilian visual lab mastology data set containing dynamic thermograms, applying components analysis resulted in three corresponding images: 1. Minimum change as a function of applied temperature or time (suggests correlation with the cancer generated heat), 2. Moderate temperature dependence (suggests correlation with veins affected by vasomodulation) and 3. Complex time behavior (suggests correlation with heat absorption due to high tumor perfusion). All components appear clear and distinct.

**Conclusions:** Applying signal processing methods to the dynamic infrared data, we found three distinct components with correspondence to understood physiologic processes. The two cases shown are self-evident of the capability of the method but are lacking supporting ground truth that is unavailable with such a limited data set. Validation of this proposed paradigm and studying furthering clinical applications has potential to create significant achievement for IR modality in diagnostic imaging.

## 1 Introduction

### 1.1 Breast cancer screening

Breast cancer is the most diagnosed cancer among women. Early detection is thought to be crucial to improve outcomes. The most common clinically used modalities for screening and diagnostic imaging of breast carcinomas are mammography, ultrasound and MRI. Mammography is regarded as the gold standard for screening and guidance of tissue sampling for suspected neoplasms. However, certain limitations exist: decreased sensitivity in dense breast tissue (more common among younger women), ionizing radiation dose to the patient (with modest expectation to result in new cancers among screening population), and inhibitive cost for use in developing, low-income countries.

### 1.2 Breast thermography

Thermography is an emission scan of medium or long wave infrared (IR) electromagnetic radiation emitted via the body, dependent upon the temperature. Thus, the IR image of the body is equivalent to an overall map of external temperature contained from individual pixels. While thermography was approved by the FDA as an adjunct method to mammography in 1982, its’ utility is very limited due to low sensitivity and poor ability to distinguish between healthy and diseased tissue. Characteristically, cancer is differentiated by an increased rate of metabolism (imaged directly in FDG PET imaging) and this may be at least partially responsible for the increase in temperature. Additionally, carcinomas are characteristic for increased blood supply via angiogenesis (used in contrast imaging techniques, via ‘enhancement’). This increase of blood flow to/surrounding the cancerous site is associated with increased ability of tissues to absorb heat. The bio heat transfer equation describing thermal propagation is given by^1^. In vivo measurements establishing tumor hyperthermia were established by Gautherie^3^, observing typical 1.5°C temperature difference between the tissue and the average temperature of the blood or similarly the temperature difference between the incoming blood to the outgoing blood. It is a consensus among all the modeling based on the bioheat equation^3,4,5^ that the large majority of the heat generate by tumor is absorb by the perfusion term. None of the models account what happens to this heat when it carries away by the veins. As we will show this heat will dominate the appearance of the diseased breast. As a result, the heat signature will appear away from the cancerous tissue where the blood flows superficially towards the skin surface, where it is dissipated to the surrounding environment.

Attempts to use thermography as a signature of cancer has limited success. The temperature of the blood pool within the arterial vascular system defines the body core temperature, assumedly unform distribution throughout the body. Thus, we can detect areas of higher heat signal than the core temperature revealed as contrast within the image, most evident as ‘hot veins’ carrying heat from high metabolic activity area. There are no publications to show evidence of correspondence between heat signature localization and anatomical tumor location. Use of IR imaging to identify neovascularization has some utility. Novel thermal imaging techniques use what is called “thermal challenge”, “cold-stress” or “dynamic imaging” where either the breast or some other part of the body is cooled while taking the thermogram, limited success was demonstrated by Y Ohashi and I Uchida^5^ who subtract the later cold series images from initial images taken prior to cold stress. Later studies^6^ failed to see an advantage in cold challenge using visual interpretation.

The authors^8,9^ in previous work demonstrated use of equivalent wave field transform (EWFT) applied to dynamic data to detect increased perfusion associated with the tumor. EWFT was developed as nondestructive testing tool and has shown success in solving the ill-posed, inverse problem associated with heat transfer. EWFT mathematically can be described as a linear transform which converts thermal diffusive propagation into a wave like propagation. The work showing heat absorbing, high perfusion regions can act as a high contrast, negative total reflector.

Some success in interpreting breast thermograms was achieved lately using artificial intelligence. Bardia^9^ at al used principal component analysis (PCA) applied to dynamic data to identify the veins as preprocessing for AI interpretation. The results are very similar to direct application of AI to the raw IR images. It appears that AI detection is based on observing neovascularization. A major disadvantage of any methods based on neovascularization is its inability to locate the cancer but only indicates existence of cancer which requires additional testing to localize it.

## 2 Method

### 2.1 Goal

The goal is to identify cancer by mapping the temperature of the surface of the breast as a response to external temperature change and identify the signature of the cancer. The problem is that most of the internal heat is carried away by the veins, thus the true location of the heat source is obscured by heat dissipating through the venous system close to the skin. By identifying heat associate with the venous and removing it we are left with heat produced by the tumor.

### 2.2 Approach

To separate the indirect heat of the venous from the direct heat of the tumor we take advantage of vasomodulation of the veins. When applying external temperature change, in order to maintain constant body core temperature, the veins constrict when cold and expand when warm. On the other hand, the metabolic rate of the tumor is expected to stay unchanged. Using component analysis, we can distinguish between the two processes. In this analysis we used sequence of thermal images of the patient breast following local temperature change using a fan. Thermal data was recorded and then analyze using PCA and independent component analysis (ICA). PCA and ICA are somewhat the opposite of each other. PCA is a technique which convert the data into sequence of orthogonal set while maximizing the variance of each set. It is mainly used to reduce the dimensionality of the data set. ICA is used to separate the data which comprised of mixed individual channels back into the unmixed sets by minimization of mutual information and maximization of non-Gaussianity. We can understand the role of PCA and ICA using the following example: We record a video of a blinking face. The first PCA image is of the face, which is common to all the images, with a superposition of the eyes. The following components are different superpositions of the eyes. If we reanalyze those images using ICA, we will recover the original individual eye position. In that case ICA undo the mixing. Both PCA and ICA has been applied in analysis of transient thermal imaging^11^ used in nondestructive evaluation (NDE). In NDE a thermal transient reflection, quite similar to our experiment, is recorded and analyzed. In that case both PCA and ICA result in very similar images of the defects. It is common to perform PCA prior to ICA, usually the low order PCA components contain most of the information while the high order PCA components are mostly noise and can be ignored while reducing the computational load of the ICA. The process we used was as follow: First we perform PCA, the first component is separated and set aside for later analysis. The next seven PCA components were analyzed, extracting four ICA components for later analysis.

## 3. Experimental Data

Data from the Brazilian Database for Breast Research^10^ with infrared images was obtained by publicly accessible methods. The protocol used for collecting the data: Room temperature was maintained between 20°C to 22°C. A fan was used to cool the skin surface of the breast until it reached 32.5°C, but for no longer than 5 minutes. The data was sampled every 15 seconds for 5 minutes. This is data of opportunity i.e., the analysis technique wasn’t in mind when the data was collected. Dynamic images were collected only from frontal view while the tumor might have been visible only from a different view. A bigger limitation was lack of medical information, patients were only classified as “sick” or “healthy” with almost no additional support information. Large number of the patient had no mammography or biopsy while still declared sick. For this publication we present two patient, patient #274 and patient #282, both left breasts. Both patients are categorized as “sick”, patient #274 had biopsy on the left breast. Patient #282 had additional information that the tumor is on the lower inner quadrant (QIM). The reason we chose those two patients is the distinct output which are very clear and speaks for themself as we lack supporting medical information.

## 4 Analysis

In order to compensate for patient motion, MATLAB *imregister* function with similarity transform followed by affine transform was applied. It followed by MATLAB *imregdemons*. *Imregdemons* is registration transformation which allows for local distortion. The *‘AccumulatedFieldSmoothing’* parameter was set between 3 and 9, which is outside the recommended value by the package. We register between adjacent images as they are quite similar. We iterate the process few times. We separated the images into left and right breasts and subtract the mean temperature of each set. As all of the following analysis steps are performed based on individual pixels, we converted the 2D image into 1D vector while keeping tab on the correspondence between the pixel’s 2D indices and the 1D vector indices. The individual image vectors are staked together to produce a matrix. We included the static image as the last frame. We kept two sets of data: a masked version of the images where only pixels from regions of interest are included and the full image. Segmentation was done manually. The transform was calculated using the masked data and applied to the unmasked data. Block diagram of the process is depicted in fig. 1.

**Figure 1:**
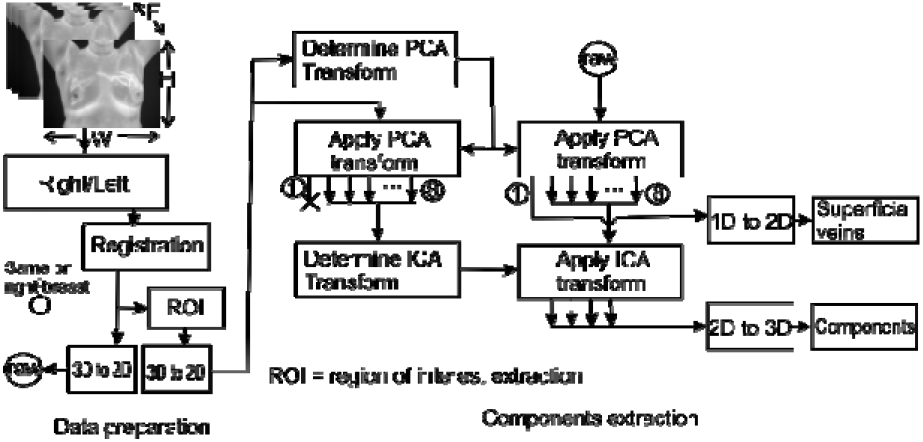
Block diagram of the analysis method.

First principal component image (e) of the veins dominates the original images, this component has the highest temperature range and the largest number of hot pixels compared to the other component images, it is very similar to the original individual frames. For that reason, we didn’t show those original images. The amplitude of each individual components (a) to (e) in the original frames are plotted in fig. 3 and 5. We scaled temperature of the displayed images based on their amplitude at the last frame. The separation between the perceived featured such as veins is very clear among the different images.

## 5 Discussion

distinguish between direct tumor heat and secondary venous heat based on vasomodulation. It is difficult to make a definite conclusion as the medical information was insufficient, but the results speak for themselves. Because of those limitation we regard this work as proof of principal rather than a thorough study. Images 1(e) and 3(e) which are first principal component are indistinguishable from the row data which are the images of the veins dispersing the high metabolic cancerous heat. The amplitude as function of frame number is plotted in fig. 3(e) and 5(e). They are consistent with rewarming and dilation of the veins. After removing the veins contribution and applying independent component analysis we obtained three clear images. Traces 3(c) and 5(d) are very similar to traces 3(e) and 5(e) the superficial veins but with longer delay to saturation, we interpretate them as deeper veins. Traces 3(a) and 5(b) change little between the frames. We interpretate them as the desired direct tumor heat. Traces 3(b), 3(d), 5(a) and 5(c) are not as clear. We assume they are reflection from heat sinks associated with high perfusion of the tumor. Such effect was observed earlier on the same data by the authors7,8, those reflections might be a combination with extra vasomodulation of the blood vessels. Both fig. 2(e) and 4(e) indicates that recording started after the rewarming was underway. Response time is composed of fan ramp up and down, heat propagation in and out and vasomodulation responses. The observed response time of fig. 2(e) is consistent with the estimated rump down of the fan^8^. Fig. 5(e) indicate deeper veins both by the time delay and by the diffused shape of image in fig. 4(e).

**Figure 2:**
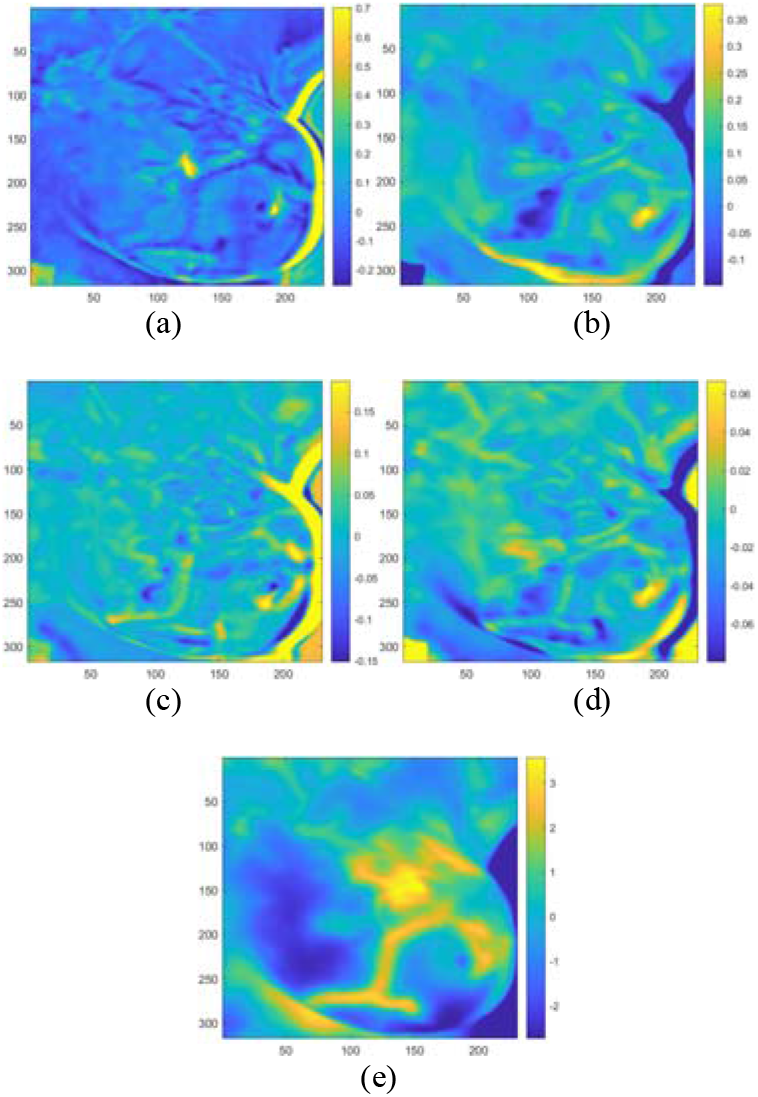
Component analysis of patient #274. (a) to (d) are independent components images, (e) is the first PCA image. Temperature in Celsius on the right is normalize to frame 20. The highest temperature is on (e) which we attribute to the veins, second highest is (a) which we attribute to the metabolic activity.

**Figure 3:**
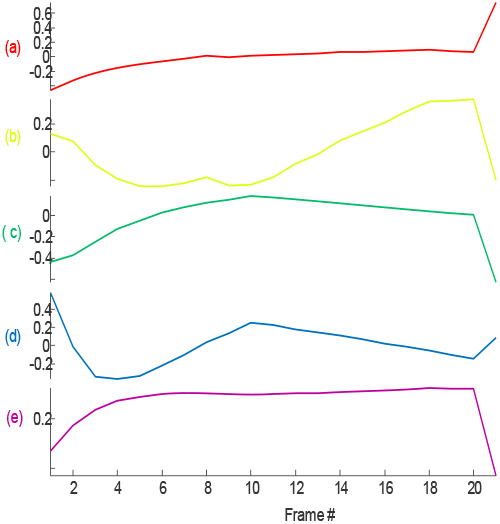
The amplitude contribution of each component (a) to (e) of patient #274 to original data frame. Last data point is of the static frame. Data is taken at 15s/frame. Trace (a) shows minimal temperature dependence, we attribute it to metabolic activity. We attribute traces (b) and (d) to thermal reflection from high perfusion. (e) and(c) are the veins response.

**Figure 4:**
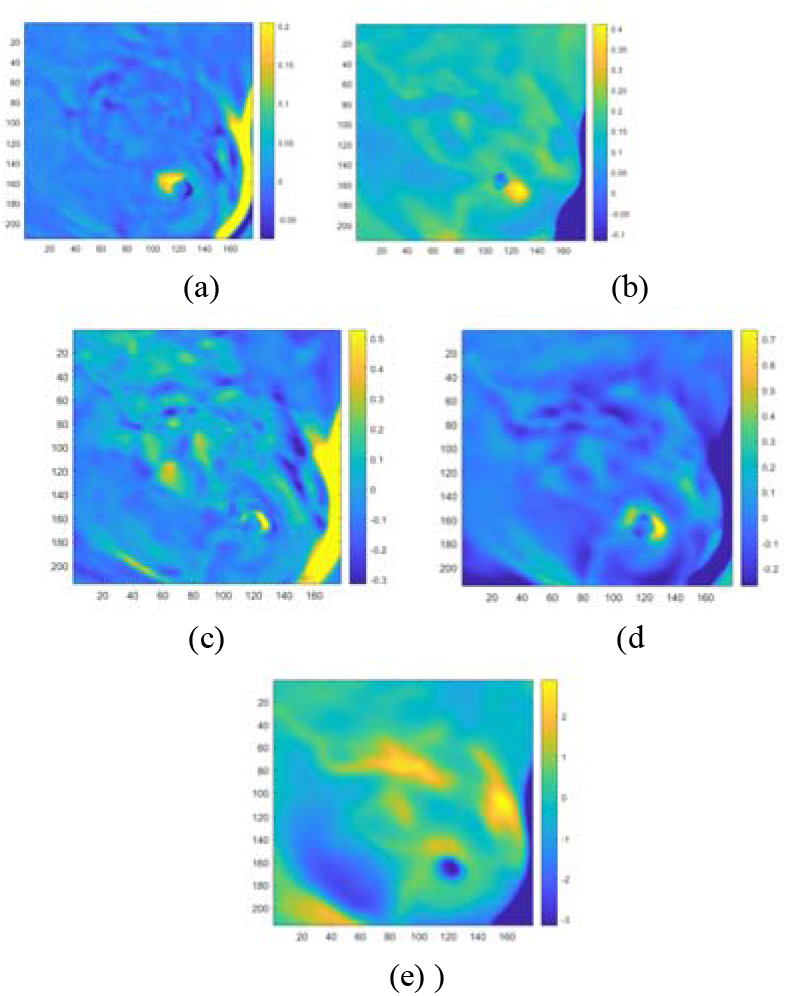
Component analysis of patient #282. (a) to (d) are independent components images, (e) is the first PCA image. Temperature in Celsius on the right is normalize to frame 20. The highest temperature is on (e) which we attribute to the veins, second highest is (b) which we attribute to the metabolic activity.

**Figure 5:**
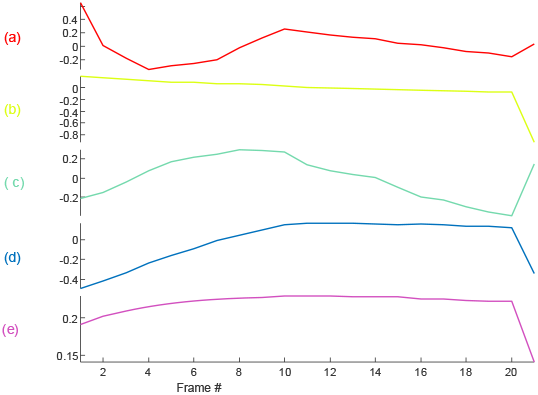
The amplitude contribution of each component (a) to (e) of patient #282 to original data frame. Last data point is of the static frame. Data is taken at 15s/frame, Trace (b) shows minimal temperature dependence, we attribute it to metabolic activity. We attribute traces (a) and (c) to thermal reflection from high perfusion. (e) and (d) are the veins response.

This work was motivated by the possibility to

## 6 Conclusion

Using data of opportunity, we have shown that employing principal and independent components analysis we can decompose sequence of dynamic breast images into components which appear to be the tumor direct heat, secondary vein heat that dominate the original image and reflected heat from high perfusion. We have to remember that this data was optimize for AI interpretation. It contains multiple static views but only single dynamic view. It is possible that the tumor may not be visualized in a single view. For the AI analysis, grouping into “sick” or “healthy” for the algorithm was sufficient, however, for our method which attempts to localize the tumor it requires correlation to tumor size and anatomical location. While the direct heat from the tumor may have been regarded as a promising objective of breast thermography, the unprocessed thermography images reveal affected veins superficially as they transport excess heat towards the skin. We have seen lately a flurry of breast thermal modeling with improved geometrical details but does not address this central issue, dominant signal of the veins which obscure the image in detection of cancer signature. Without processing to remove such dominant artifact, detecting the cancer signal directly is likely impossible, reflective of the limited progress this modality has achieved. In this work, we showed that based on vasomodulation one can differentiate between signals, suggested to represent the desired heat of the tumor and the secondary heat from the veins. Both the images and their time dependence are consistent with our expectations. Additionally, reflected heat resulted from the high perfusion of the tumor was also identified.

## Acknowledgment

I would like to thank Dr. Mark Mandelkern and Dr. Paul F. Schippnick for critical reading and suggestions.

**Meir Gershenson** is a retired physicist. He received his BS and PhD degrees in physics from Tel Aviv university 1967. and 1976, respectively. His PhD is in superconducting materials. He worked at Sperry corporation on superconducting electronics and the US Navy in mine countermeasure. He is the author of more than 30 journal papers and 15 patents. His current research interest is imaging through diffusion processes.

**Jonathan Gershenson** is an attending physician of Nuclear Medicine at the VA Northern California Healthcare system. He graduated from medical school at Kansas City University of Medicine & Biosciences in 2016 before completing residency in Nuclear Medicine at UCLA/VA West Los Angeles Medical center in 2021. He has additional training in medical research methodology through the UCLA Translation Science certificate program.

